# Honeybee hive density has consequences for foraging bumblebees in Irish heathlands

**DOI:** 10.1101/2025.03.07.642004

**Authors:** Katherine L.W. Burns, Lina Herbertsson, Dara A. Stanley

## Abstract

Heathlands provide vital foraging resources for wild bumblebees, as well as managed honeybees that are brought to heathlands in late summer for heather honey production. With this increased honeybee activity, there is potential for competition for floral resources between honeybees and bumblebees. We studied how increasing number of honeybee hives in Irish upland heathlands, influenced honeybee abundance, nectar availability and bumblebee abundance, size and activity at two distances from the honeybee hives. In general, an increasing number of hives resulted in elevated honeybee abundance, and - while we observed no change in nectar volumes - an increase in flower-visitation activity (flowers/minute) by individual bumblebee workers. Furthermore, bumblebee worker size declined with hive density, suggesting that larger workers - which have a longer foraging range - may have foraged further from the hives, or possibly that smaller workers may have been forced participate in foraging to compensate for smaller rewards. This highlights the need to consider interactions between wild and managed bees for floral resources in land management and conservation to ensure the protection of wild bumblebee populations and the continued productive management of honeybee populations for profitable honey production.

## Introduction

Wild bee populations are in decline due to a variety of anthropogenic pressures, including discontinuous forage availability and spread of diseases (1–3). Because honeybees and wild bees exploit similar floral resources (4–6) and honeybees can transfer pathogens to wild bees (7, 8), the increasingly widespread use of managed honeybees (*Apis mellifera*) (9) has the potential to exacerbate these threats. Indeed, beekeeping can reduce the availability of floral resources with consequences for wild bees (e.g. 10, 11). Although there is still no clear link between beekeeping and negative population development in wild bees (12), there is evidence that beekeeping or altered levels of honeybees can reduce local wild bee abundance (e.g. 10, 13), alter realized foraging niches (14) and change pollen foraging efforts (15). Altered levels of honeybees have also been shown toreduce bumblebee forager size (16), potentially as a result of food limitation during larval development (17, 18), altered colony labour division - with a higher proportion of small individuals leaving the nest - to compensate for low foraging efficiency (19), or larger sized worker individuals – with a longer maximum foraging radius (20) – escaping competition by foraging at sites with lower honeybee densities. Outside of their native range, honeybees have also been observed to influence wild bee reproduction negatively (15).

Increasing awareness about potentially negative effects of honeybees on wild bees has driven an interest to protect wild bees from competition, also in Europe (21), where honeybees are native or have existed for thousands of years (22). Authorities in many European countries are working actively with beekeepers to discuss solutions, or have even banned commercial beekeeping in protected areas (21). However, such bans have been criticized for being inefficient due to the long foraging ranges of honeybees, and for causing conflicts between beekeepers and conservationists (23). To effectively safeguard wild bees without unnecessary restrictions for beekeepers, it is crucial to understand when and where interactions occur and could become a competitive problem for wild bees.

As of 2022, 66% of studies measuring interactions between managed and wild bees reported negative effects on the wild bees (12). Compiled evidence indicates that competition occurs mainly when bee populations are limited by floral resources (13, 24–26), and - although extreme honeybee densities can influence wild bees negatively in flower-rich environments (27–29) - this suggests that negative consequences of honeybees primarily occur in environments with limited flower availability.

European heathlands are flower-rich environments, providing habitats and forage for a variety of insect species (Moquet et al., 2017b, Baude et al., 2016, Descamps et al., 2015, Forup et al. 2008, Mahy et al., 1998). In the late summer, when other floral resources are scarce, insect pollinators, particularly wild bumblebees, depend on the late-flowering, nutrient-rich heather species, such as ling heather (*Calluna vulgaris*), that grow in these habitats (30, 31). In addition, evidence has shown that the nectar of *C. vulgaris* also acts as a “bee medicine,” preventing parasitic infections in bumblebees (32). However, not only are these heathlands important for bumblebees, they also constitute an economically valuable resource in terms of honey production (31, 33). Recent findings have shown that the antioxidant capacity and corresponding human health benefits of *C. vulgaris* honey may be comparable to that of Manuka honey, which could increase the value and demand for heather honey (33), incentivising beekeepers to expand heather honey production.

This raises the question whether the floral resources in heathlands are sufficient to prevent competition among bee species. Previous studies assessing correlations between foraging bumblebees and honeybees in heathlands found weak evidence for competition (26, 34), but because bumblebees as well as honeybees are attracted to, and benefit from, flowering heather, such methods may be misleading. To understand if honeybees pose a threat to wild bumblebees in heathlands, it is necessary to design a study based on the nearness to, or density of, hives rather than observations of honeybees.

In a full-factorial design, we aimed to determine the effect of distance to (250 m and 1000 m) and number of (a gradient of 0-35 hives) honeybee hives on (1) honeybee abundance, (2) resource availability, and (3) wild bumblebees, more specifically (3a) foraging activity (flowers/minute), (3b) size, and (3c) abundance. We expected increasing hive numbers and decreasing distance to the hives to (1) increase honeybee abundance, (2) with negative consequences for nectar volumes. Because the length of flower visits increases with the availability of floral rewards (35–37), we expected (3a) increased flower-visitation activity (flowers/minute) and due to reduced resource availability, we expected (3b) smaller size and (3c) lower abundance of bumblebee foragers.

## Materials and Methods

### Site selection

The Dublin and Wicklow Mountains (53.141 N, −6.311 W) in eastern Ireland are dominated by heath (nearly 40%; 12,784 ha), blanket bog, and grassland habitats, and are commonly used for sheep-grazing, peat cutting, and forestry (38). Large areas of the landscape are protected within Wicklow Mountains National Park, most of which also overlaps with Natura 2000 designated sites (38, 39). In the late summer, some beekeepers bring their most productive hives to the mountains to produce single-origin heather honey (Irish beekeepers, personal communication). Currently, beekeepers are not allowed to place their hives within the Wicklow Mountains National Park boundaries (Wicklow Mountains National Park, personal communication).

Seven independent apiaries, surrounded by a minimum 25% heathland cover within a 3000 m radius (maximum cover was 83%), were identified, each containing between 3 and 35 hives. Additionally, we selected one area without any known apiaries within 3000 m (number of hives = 0). Sites were chosen based on the locations of seasonal apiaries (which were mapped as comprehensively as possible) to ensure each apiary was at least 1000 m away from the next. Sites were also selected based on the percentage of heathland cover in the area surrounding the apiaries, determined using the 2018 CORINE land cover dataset (40), to ensure that the surrounding landscapes of the sites were similar in terms of composition and the availability of floral resources.

Within each site, two 100 m radius sampling areas, where heather was the dominant flowering species, were selected: one sampling area at 250 m from the apiary (or selected corresponding position at the control site) and another 1000 m from the apiary (Figure 2). All sixteen sampling areas (two per site) were separated by a minimum of 1000 m from any other apiary and unassociated sampling areas. Sites (Figures 1 & 2) were surveyed twice between 6^th^ August – 6^th^ September 2020 (minimum of two weeks between survey rounds). The first round of observations conducted at each site took place within a week of beekeepers bringing their hives up to the heather.

**Figure 1.**
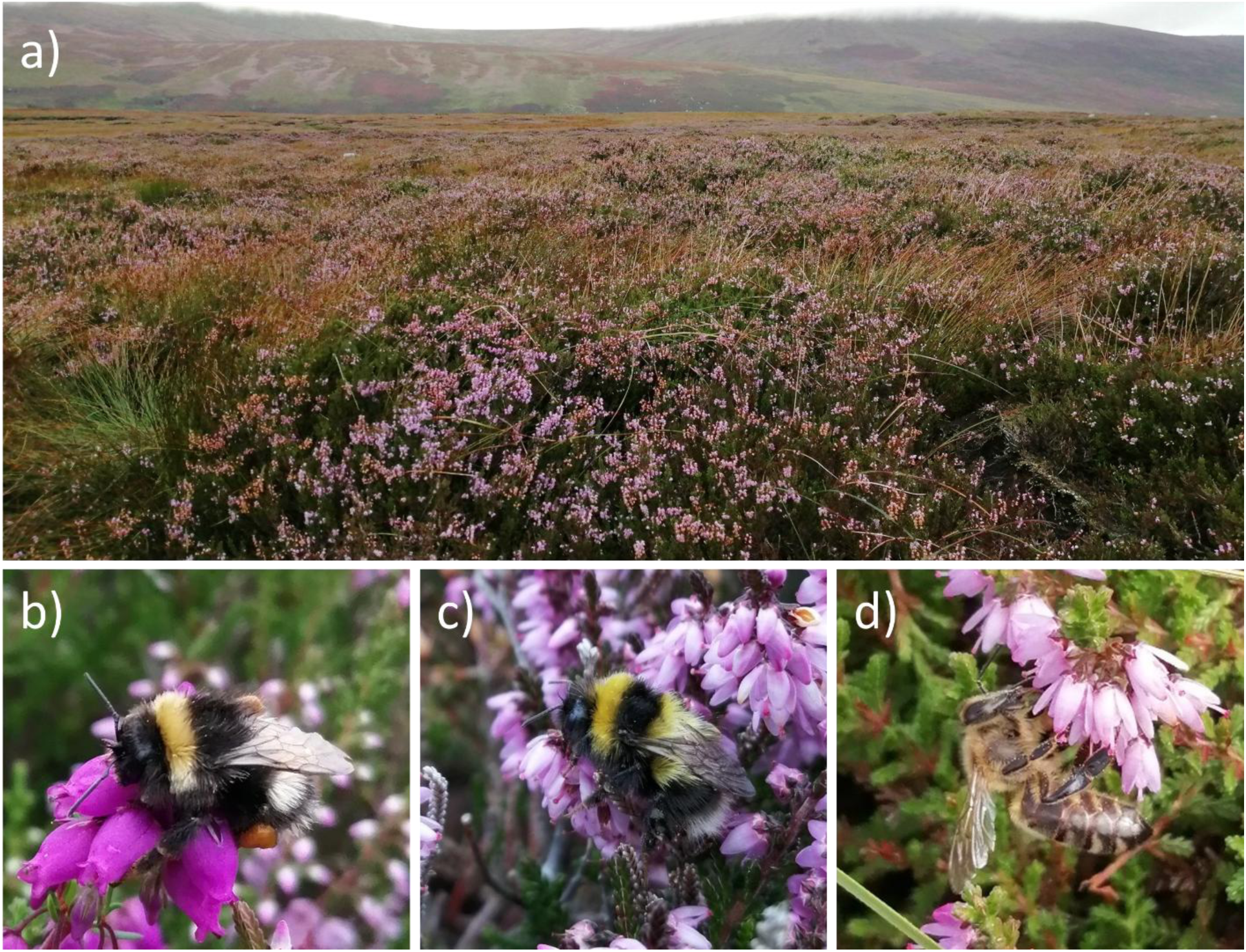
Example of a study site and bees found foraging on heather species: (a) heathland study site, (b) *Bombus lucorum agg.* on *Erica cinerea*, (c) *B. jonellus* on *Calluna vulgaris*, and (d) *Apis mellifera* on *C. vulgaris*.

**Figure 2.**
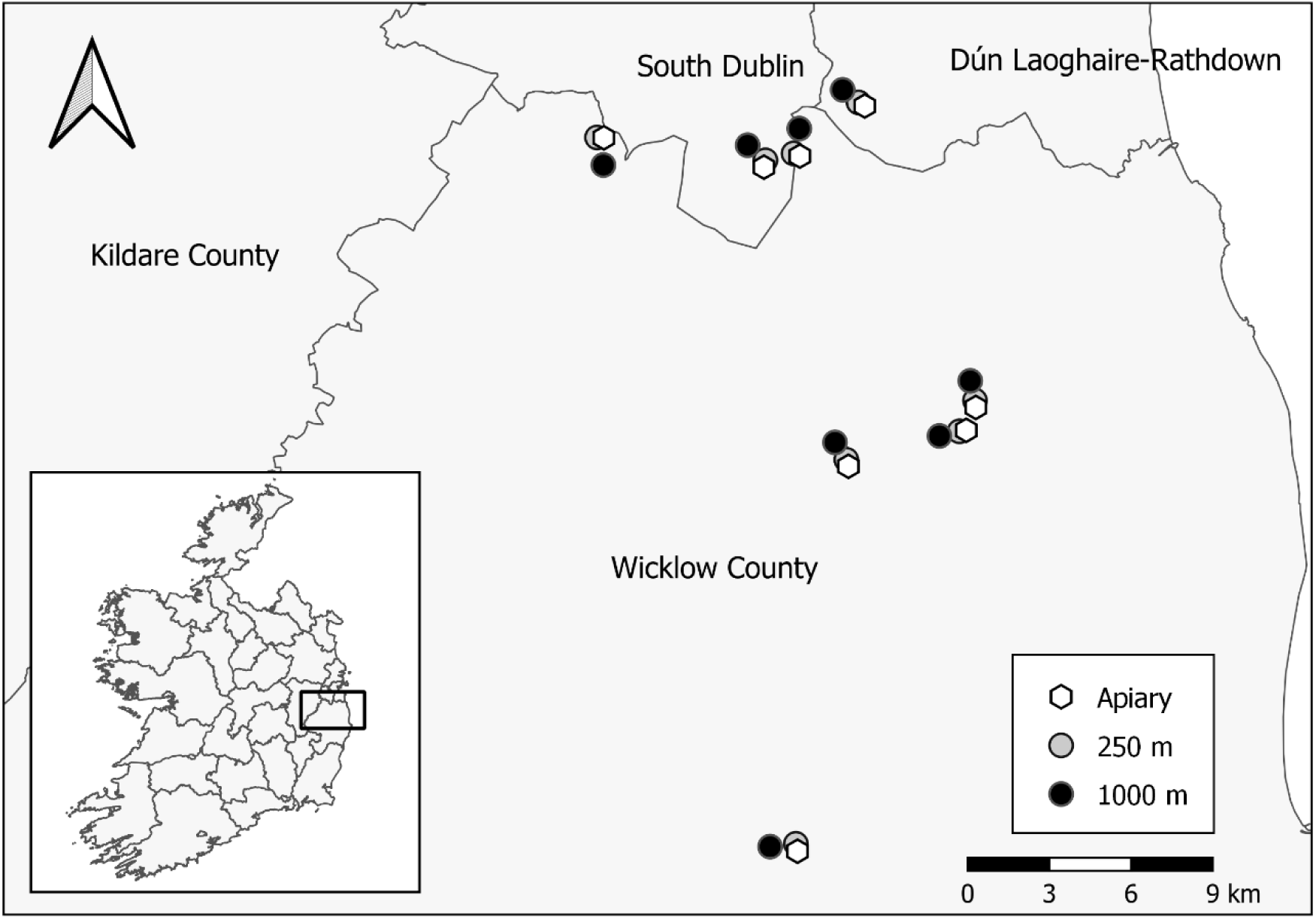
Locations of apiaries (white hexagons) and associated sampling areas: 250 m (grey dots) and 1000 m (black dots); (Map Credit: 41)

### Bee abundance and foraging activity

During each survey round, we recorded abundance of honeybees and bumblebees on two 100m transect walks per sampling area, during which we also recorded the species, caste, visited plant species and presence of pollen loads of the recorded bees. We recorded bumblebee foraging activity by following up to five *B. lucorum agg.* worker individuals - which corresponded to roughly 77% of the bumblebee individuals in our sampling areas - for one minute, or shorter if they moved out of sight, noting the number of flowers visited and the forage species. One hour was allocated to these observations and at sites where bumblebee abundance was low, we therefore sometimes followed fewer individuals. Activity was calculated as the number of flowers visited per minute. All transect walks and bee observations were carried out on warm, dry days (11 – 22° C) with low wind speeds (1 – 28 km/h).

### Bumblebee size

To measure bumblebee size, we aimed to capture at least 15 workers of each detected species during each visit to a sampling area. The sampling area was walked for one hour and collectors opportunistically caught worker bees observed foraging on heather. Bumblebee abundance and richness were low at certain sites, in which case fewer than 15 bumblebees were captured. Bees were cooled until immobile, and their intertegular distance measured using a Vernier calipers before being released. To avoid catching the same bee more than once, we kept all captured bees in a cooler until measurements were completed and then released all of the captured bees together. Bee sampling for size measurements took place after transect walks and activity observations were completed to avoid influencing bee abundance and foraging activity.

### Floral resources

To calculate the floral abundance in each sampling area, percentage cover of each heather species in full flower and partial flower was recorded in five randomly chosen 1 m^2^ quadrats on each survey round. Total available percent cover of flowering heather foraging resources was calculated according to methods outlined by Franklin et al. (26).

To assess nectar availability, nectar standing crop was measured at the end of the day during each site visit. Later, we decided to exclude nectar samples from the first visit to one of the sites, where it rained before nectar was collected. Nectar was sampled from 10 open flowers from each heather species present in the sampling area, and the volume was measured to the nearest microlitre using 0.5 microlitre microcapillary tubes.

## Data Analysis

### General procedure

All data were analysed in R version 4.1.1 (42). We used (generalized) linear mixed effects models (package “glmmTMB”, 43) with site as random factor to account for repeated sampling in time and space. When there was more than one data point per distance (250 or 1000 m), we nested distance within site to account for non-independence of data points from the same sampling area. We specified hive density (0-35), distance from the hives (250 or 1000 m), the interaction between these two (hive density × distance), and sampling round as fixed factors. We evaluated residuals using Kolmogorov-Smirnov, outlier and dispersion test, as well as quantile plots from the DHARMa package (44) and verified that the random effects were estimated and positive. We interpreted p values with a type two ANOVA, allowing interpretation of main term effects without the removal of interactions, and consequently we retained all variables in the model, including non-significant interactions. We interpreted interactions with contrasts (joint_tests from the package emmeans; 45) and from back-transformed model predictions with 95% confidence intervals (predict.glmmTMB from the package “glmmTMB”, 43).

### Specific model details

To evaluate the influence of our model design on abundance of honeybees and bumblebees, respectively, we initially specified a model with poisson distribution. To avoid zero-inflation, we aggregated data per site, distance and round. Because of strong under-dispersion, we added 1 to the number of honeybees, after which we performed a log-transformation and specified a gaussian distribution. We evaluated the effect on worker thorax width and activity (flowers per minute and individual) of *B. lucorum agg.*, specifying gaussian distribution. Because *B. lucorum agg.* and *B. jonellus* together accounted for 93% of the collected data on thorax width (104 *B. jonellus*, 205 *B. lucorum*), we excluded data from other species (1 *B. lapidarius*, 4 *B. monticola*, 17 *B. pascuorum*) and specified bumblebee species (*B. lucorum agg.* or *B. jonellus*) as a fixed factor in the model. As the raw datasheet for a subset of thorax results went missing and these data could not be double checked before submission, we ran analyses both with and without data from this sheet. When analysing activity (flowers per minute) of individual *B. lucorum agg.,* we removed two bees that had visited a mixture of plant species and added genus of the visited species (*Calluna* 89*%, Erica* 11%) as a fixed factor to account for shorter handling time of *Calluna*. To understand if honeybees influenced nectar availability of open flowers while accounting for site specific variations in nectar production (estimated from bagged flowers), we added treatment to the interaction, resulting in a three-way interaction between treatment (open or bagged), distance and hive density (treatment × hive density × distance). We square root transformed volume to normalize the residuals. To improve the residuals further and account for a strong collector bias, we specified collector identity as a fixed factor. We initially included data from all three heather species and specified genus, or species, as a fixed factor to account for differences between *Calluna* and *Erica*, or among the three species. However, these models suffered from highly significant Kolmogorov-Smirnov and outlier tests and had strong quantile deviations. Therefore, we only included *C. vulgaris* - which dominated the floral cover in all sampling areas and accounted for more than 65% of the nectar samples. We verified that the proportional cover of flowering heather was unrelated to our study design specified a model with gaussian distribution. With post-study access to more detailed land use data (46) than provided by CORINE (EPA 2018), which we used for the design, we specified a model with gaussian distribution and without distance in the random structure to test if the proportion of heathland was evenly distributed across our variables of interest. Because this model showed that the proportion of heathland was lower at the distant sampling areas, we ran an additional set of models, where we for each of the previously described models replaced distance to the hives with proportion of heathland (Text S1).

## Results

### Bee abundance

A total of 182 honeybees and 145 bumblebees (Table S1) were observed during transect observations, the majority of which were foraging on *C. vulgaris* (Table S2). Of the bumblebee and honeybee workers observed, only a minority were carrying pollen (34% and 13%, respectively).

Honeybee abundance was unrelated to the interaction (n = 32, χ^2^_1_ = 0.94, p = 0.33), increased with hive density (n = 32, χ^2^_1_= 3.90, p = 0.05, Figure 3a), was lower during the second round (n = 32, χ^2^_1_ = 5.06, p = 0.02, Figure 3b) and unrelated to distance (n = 32, χ^2^_1_ = 1.82, p = 0.18, Figure 3a). Bumblebee abundance was marginally related to an interaction between hive density and distance from the hives (n = 32, χ^2^_1_ = 3.13, p = 0.08, Figure 4a). Bumblebee abundance was higher during the second round (n = 32, χ^2^_1_ = 28.50, p < 0.0001, Figure 4b).

**Fig. 3.**
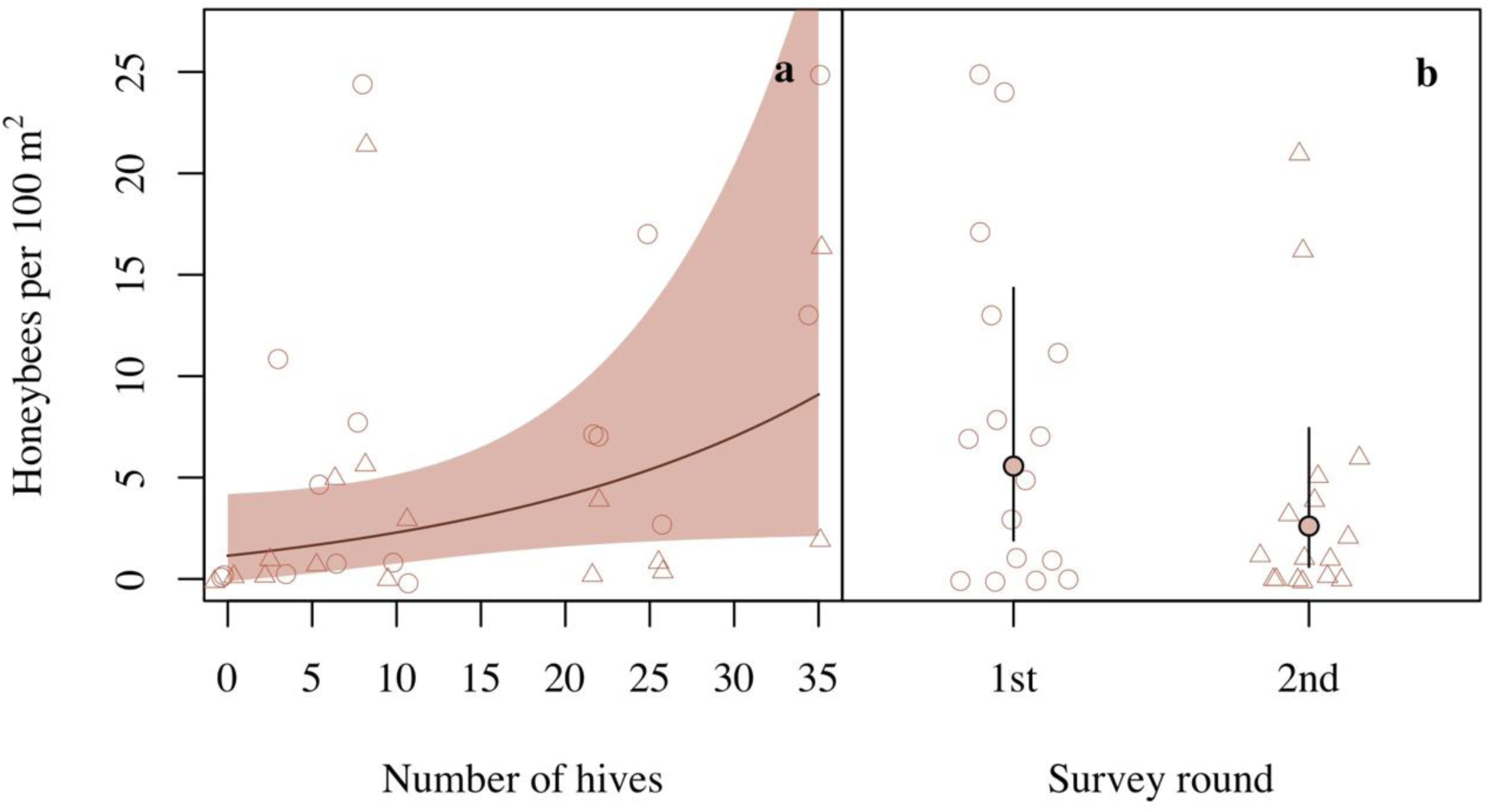
**a)** Honeybee abundance increased with hive density b) and was lower during the second survey round. Figures display back-transformed predictions with 95% confidence intervals. Open circles and triangles display raw data from the first (circles) and second (triangles) survey round. Raw data points have been jittered to reduce overlap and increase their visibility.

**Figure 4.**
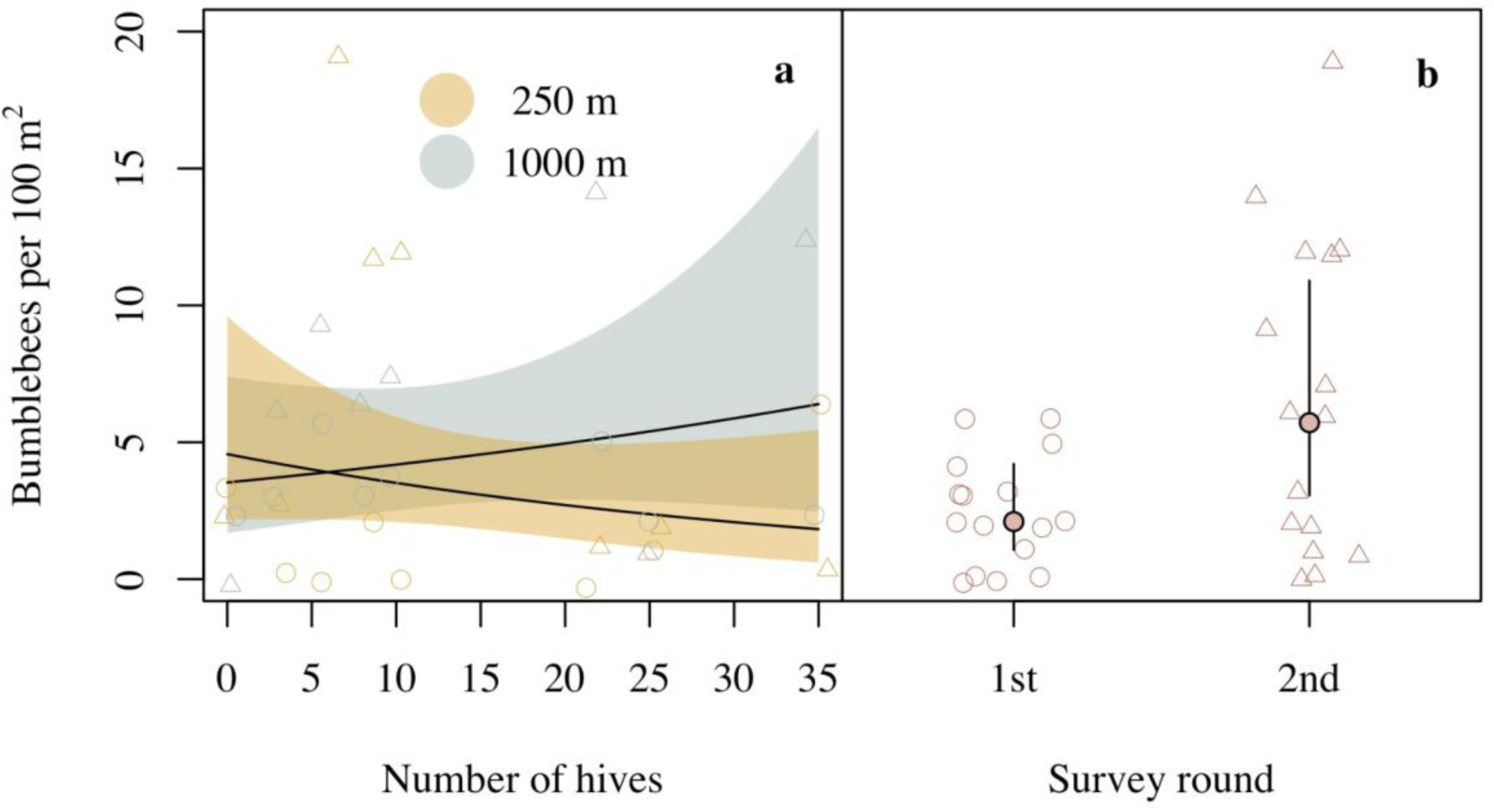
**a)** Bumblebee abundance was related to a marginally significant interaction between hive density and distance to the hives and **b)** was higher during the second survey round. The figure shows back-transformed model predictions and 95% confidence intervals. Open circles and triangles display raw data from the first (circles) and second (triangles) survey round. Raw data points have been jittered to reduce overlap and increase their visibility.

### Flower visitation and size of bumblebee workers

The number of flower visits per minute by an individual of *B. lucorum agg.*was unrelated to the interaction (n = 131, χ^2^_1_ = 0.01, p = 0.93), distance (n = 131, χ^2^_1_ = 0.12, p = 0.73) and round (n = 131, χ^2^_1_ = 0.02, p = 0.89), but increased with hive density (n = 131, χ^2^_1_ = 4.49, p = 0.03, Figure 5a) and was higher for bees visiting *Calluna* than *Erica* (n = 131, χ^2^_1_ = 18.06, p < 0.0001, Figure 5b). Thorax width of *B. lucorum agg.* and *B. jonellus* was unrelated to the interaction (n = 309, χ^2^ < 0.01, p = 0.99) and distance (n = 309, χ^2^ = 0.55, p = 0.46), but declined with hive density (n = 309, χ^2^_1_ = 7.27, p = 0.007, Figure 6a). Thorax width was smaller during the second round (n = 309, χ^2^_1_ = 36.49, p < 0.001, Figure 6b) and *B. jonellus* individuals were smaller than *B. lucorum agg.* (n = 309, χ^2^ = 127.28, p < 0.001, Figure 6c). These results were consistent also without the data from the missing data sheet (n = 275, hive density ⨉ distance: χ^2^_1_ = 1.72, p = 0.19, hive density: χ^2^ = 5.36, p = 0.02, distance: χ^2^ < 0.01, p = 0.96, round: χ^2^ = 47.77, p < 0.0001, species: χ^2^_1_ = 147.46, p < 0.0001).

**Figure 5.**
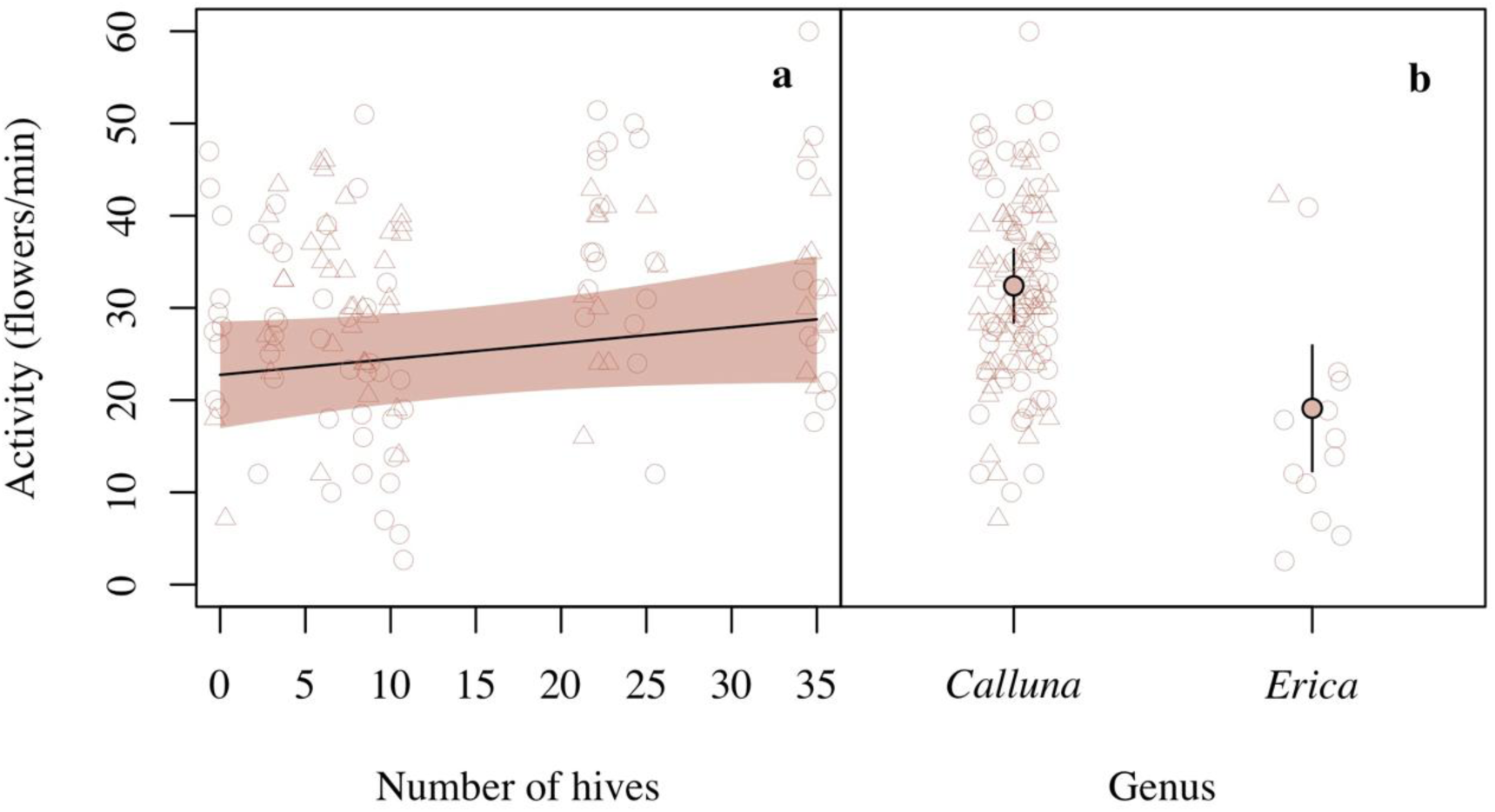
Activity (flower per minute) of individual *Bombus lucorum agg.* workers **a)** increased with hive density and **b)** was lower for individuals visiting *Erica sp.* than for bees visiting *Calluna vulgaris*. The figure shows model predictions with 95% confidence intervals. Raw data are displayed as circles (first survey round) and triangles (second survey round). Raw data points have been jittered to reduce overlap and increase their visibility.

**Figure 6.**
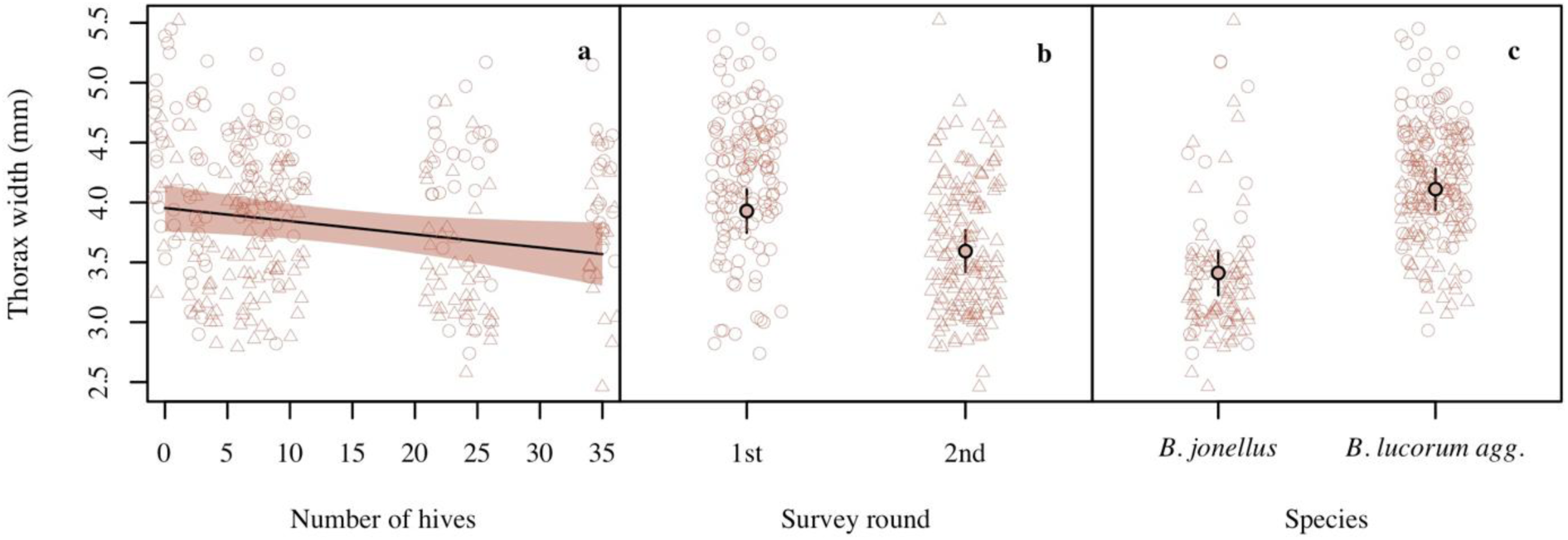
**a)** Thorax width of bumblebee workers declined with hive density, **b)** workers were larger during the first than the second survey round and **c)** workers of *B. lucorum agg.* were larger than those of *B. jonellus*. Model predictions and 95% confidence intervals are shown a) as a line and shaded areas, and b-c) as filled circles and whiskers. Raw data are displayed as open circles (first survey round) and triangles (second survey round). Raw data points have been jittered to reduce overlap and increase their visibility.

### Flower resources

The most common floral resource was flowering ling heather (*C. vulgaris*, mean ± sd = 34 ± 24%), followed by bell heather (*E. cinerea*, mean ± sd = 1 ± 2%) and cross-leaved heather (*E. tetralix*, mean ± sd = <1 ± 3%). Total available percent cover of flowering heathers within each transect was unrelated to hive density (n = 160, χ^2^_1_ = 0.83, p = 0.36), distance from them (n = 160, χ^2^ = 0.65, p = 0.42) and their interaction (hive density ⨉ distance: n = 160, χ^2^ = 1.30, p = 0.25), but declined from the first to the second round of observation (n = 160, χ^2^ = 47.90, p < 0.0001). Nectar volumes were larger in bagged flowers (n = 620, χ^2^_1_ = 70.91, p < 0.0001) and during the first round (n = 620, χ^2^_1_ = 42.55, p < 0.0001), and differed between the two collectors (n = 620, χ^2^_1_ = 20.60, p < 0.0001) but were unrelated to honeybees because treatment (bagged or open) did not influence any of the honeybee-related variables (n = 620, treatment ✕ hive density ✕ distance to hives: χ^2^_1_ = 0.16, p = 0.69, treatment ✕ hive density: χ^2^ = 0.54, p = 0.46, treatment ✕ distance: χ^2^_1_ = 2.42, p = 0.12, Figure S1), and hive density (χ^2^_1_ = 0.00, p > 0.99), distance (χ^2^_1_ = 0.70, p = 0.40) and their interaction (χ^2^_1_ = 2.02, p = 0.15) also had no influence on nectar volumes. We failed to extract any nectar at all from 31% of the open flowers, and from 5% of the bagged flowers.

### Land use

The proportion of heathland within 1000 m varied between 26 and 93% and was unrelated to the interaction (n = 16, χ^2^_1_ = 1.33, p = 0.25) and hive density (n = 16, χ^2^_1_ = 0.33, p = 0.57), but was generally lower around the 250 m sampling area than around the 1000 m sampling area (n = 16, χ^2^_1_ = 16.73, p < 0.0001). Around the 250 m sampling area, the proportion of heathland varied between 26 and 88%, whereas it varied between 40 and 93% around the 1000 m sampling area. The proportion of heathland had no influence on honeybee abundance, bumblebee worker activity or proportional flowering heather (Text S1). Bumblebee abundance decreased with increasing proportion of heathland (n = 32, χ^2^_1_ = 6.45, p = 0.04), and this effect was unrelated to hive density (heathland × hive density, n = 32, χ^2^_1_ = 0.32, p = 0.57). Thorax width was related to an interaction between hive density and proportion of heathland (n = 306, χ^2^_1_ = 6.54, p = 0.01, Figure 7), reflecting a generally positive influence of the proportion of heathland. Nectar volumes were related to a significant interaction between treatment (open or bagged), hive density and proportion of heathland (Text S1, Figure S2).

**Figure 7.**
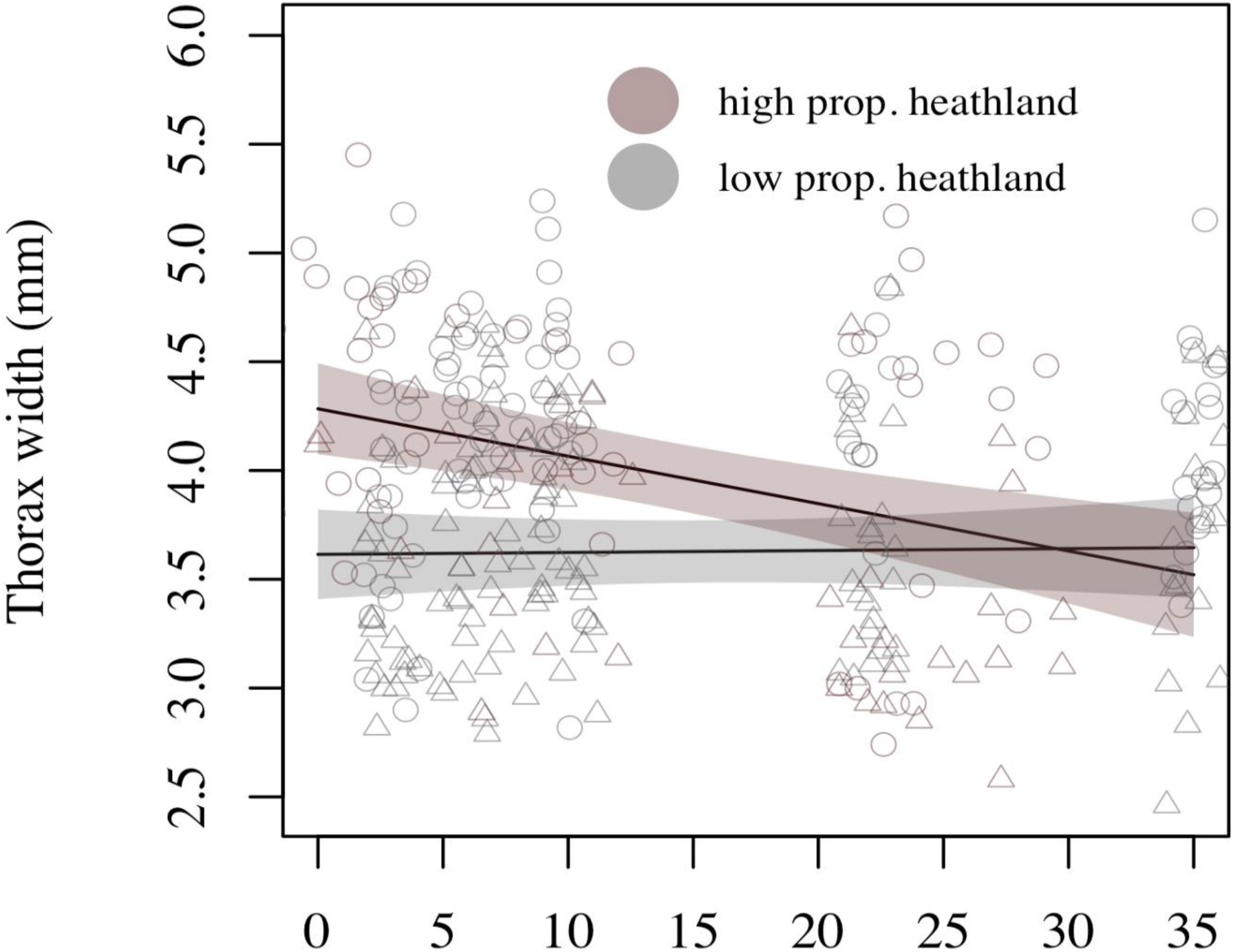
Thorax width of bumblebee workers decreased with hive density when the proportion of heathland was above average (pink), but was unrelated to hive density when the proportion of heathland was below average (grey). The figure shows model predictions

## Discussion

Previous studies assessing competition between honeybees and wild bees in European heathlands have been based on correlations between the abundances of honeybees and bumblebees, with a risk that observed patterns are driven, or obscured, by similar response to floral resources or differences in floral preference among bee species (26, 34). Using a full-factorial design, combining a gradient of honeybee hives with different distances from these hives, we provide evidence verifying that honeybees can influence bumblebees in these flower-rich systems.

### No evidence for reduced nectar but marginally reduced foraging efficiency

Heathlands are largely homogenous landscapes in terms of floral richness (47), especially in the late summer and early autumn when most flowering plant species begin to go out of bloom and *C. vulgaris* becomes the primary source of forage in the landscape (34). Not surprisingly, *C. vulgaris* - which dominated the floral cover in our sampling areas - was the most frequently visited forage species by both honeybees and bumblebees, indicating a large overlap in resource use. Previous studies have shown that in areas with high honeybee activity, nectar and pollen resources can become more rapidly depleted (10, 27). When studying nectar standing crop, we found no evidence for such effects. Although this suggests that honeybees may not have a large influence on nectar availability in this system, it may also be related to the variability and limitations of measuring and using nectar standing crop (48). It is also important to underline that our nectar sampling suffered from a few drawbacks; firstly, we did not empty flowers before the bagging and nectar of bagged flowers may thus partly reflect pre-bagging conditions. Secondly, we failed to extract nectar from 6% of the bagged flowers, suggesting a large variation unrelated to the honeybees. Thirdly, a strong collector bias revealed a difference in accuracy between data collectors, which may have influenced the results further, highlighting the need for larger sample size and balanced sampling efforts of nectar collectors in future studies.

While we found no evidence for reduced nectar availability, the number of flowers visited per minute by individuals of *B. lucorum agg.* increased with increasing hive density. Because bees spend more time on flowers with more floral resources (35–37), this pattern may reflect smaller nectar or pollen rewards, and consequently reduced forage efficiency, in areas with high hive density. The exact mechanism behind the slightly shorter visits is, however, unclear. One option is that the bees simply stayed shorter because they emptied the nectar or pollen rewards of each flower quicker, another option is that the bees shifted from pollen collection, which is time-consuming but necessary for colony development, to nectar foraging, which has direct consequences for colony maintenance. Previous observations suggest that competition with honeybees can force bumblebees to reduce their pollen collection efforts, presumably because colony maintenance is more urgent than investing in colony growth when nectar resources are scarce (15). Thus, the increase in number of flowers per minute may result either from shorter flower visits because of smaller rewards, reduced efforts to collect pollen, or a combination thereof. Regardless of the mechanisms, if occurring over long periods, reduced foraging efficiency and pollen harvest could be expected to harm fitness, unless the bumblebees manage to compensate by escaping competition or increasing their foraging efforts.

### Smaller bumblebee foragers as an indication of resource shortage

It is possible that the smaller worker size in areas with higher hive densities reflects an attempt to compensate for reduced foraging efficiency. The pupal stage of bumblebees lasts for around two weeks (18) and our first visit to the sites occurred within a week after the introduction of honeybees. Within this timespan it is impossible to observe effects from distorted colony development and reduced offspring size. Instead, the smaller size of workers in areas with high hive densities may result from changed labor division within the colony. Smaller workers are less efficient foragers (49) and are often responsible for in-hive duties (19). It has been suggested that resource scarcity due to competition with honeybees can force small workers to participate in foraging to compensate for low colony-level foraging efficiency (16), which may explain why we observed smaller foragers in areas with high hive densities.

Another potential explanation could be that larger bodied bumblebees, which have larger foraging ranges than smaller bodied bumblebees (20), escaped competition by displacing further from our study sites in response to the increased presence of honeybees, while smaller individuals remained (27). Such displacement could explain why thorax width was particularly negatively related to hive density in sampling areas surrounded by high proportion of heathland, where bees with a sufficiently large foraging range had access to alternative foraging habitat, whereas smaller bees with shorter foraging range may have been forced to remain near the hives. This may also explain why bumblebee abundance was related to a marginally significant interaction between increasing hive numbers and distance to the hives, with a slight decline 250 m from the hives and a contrasting increase 1000 m away. It is thus plausible that reduced thorax width and marginal effects on bumblebee densities reflect temporary behavioral adaptations to competition rather than inhibited colony development. Yet, we cannot exclude that the energy and extra time required to collect resources further away from the nest, or suboptimal adjustments of task partitioning resulting in less focus on brood-care, affect colony growth (50), and as a consequence, colony fitness (51).

Given that we started monitoring less than a week after the introduction of honeybee hives, inhibited larval growth due to resource depletion by honeybees is unlikely to be the only - or even the main - driver behind the smaller thorax width, but we cannot rule out that it contributed to the smaller worker size during the second round. As smaller workers collect less food per unit of time than larger workers (49, 52), a reduction in worker size may have serious implications for bumblebee colonies later in the season. For example, the body size of new queens, which is largely influenced by larval food availability (18) is strongly linked to hibernation survival rate (53, 54). Competition with honeybees can indeed reduce queen densities during the next spring, but whether this is driven by reduced queen production or hibernation survival has not been evaluated (55). Because a reduction in food provisioning resulting in smaller sized workers could affect overall colony reproductive fitness, it is crucial that future studies disentangle the full mechanisms behind the smaller worker size in areas with high honeybee densities observed in this and other studies (16) as well as their long-term consequences for bumblebee populations.

In this study, the size of bumblebee workers declined from the first to the second survey round, which contrasts with the expected development in healthy colonies, where the average bumblebee worker size increases over the course of the flight season (56–58). A reduction in flowering heather during the same period suggests that the flowering peak had passed, potentially resulting in a resource shortage, forcing the smaller workers to participate in foraging (c.f. 19). During the same period, we observed a decline in honeybee abundance and a contrasting increase in bumblebee abundance. Although it is possible that the contrasting patterns reflect that bumblebees avoided foraging in areas with high honeybee densities and returned to the sampling areas during the second round, when the honeybee numbers went down (14), another explanation could be that the two groups responded differently to resource shortage because of different foraging strategies. While the large maximum foraging range and advanced communication of honeybees may have allowed them to identify and exploit patches that were still rich in flowers (59), bumblebees may instead have compensated for low foraging efficiency by recruiting more - and thus smaller – workers (19).

### Implications for bumblebee conservation and future research

Increasing hive density resulted in higher honeybee densities, smaller bumblebee workers and an increase in flower visitation activity (flowers/minute) by bumblebee workers, and these effects were equally noticeable at the two distances from the hives. Indeed, honeybee foraging ranges can vary greatly (< 500 m – 10,000 m) in heathlands, depending on the size and quality of forage patches in the landscape (60). In Mediterranean shrublands, competition between honeybees and wild bees peaked at around 600-900 m from the hives and was still noticeable 1.2 km away, although - importantly - hive densities in the previous study were considerably higher (30.9 ± 21.8, s.d.) than in our study (15.57 ± 11.87, s.d., excl. the control site) (27). From an applied conservation perspective this is an important finding as it suggests that 1000 m is not enough to fully protect wild bumblebees from potentially competitive interactions with honeybees in Irish heathlands. It is, however, important to acknowledge that competition for food is a common ecological phenomenon among co-occurring species with overlapping niches and the sole observation that wild bee species change their behavior in relation to honeybees does not imply that their fitness is reduced (5). Although we observed that honeybees induced changed foraging patterns in wild bumblebees, the consequences for bumblebee fitness remain unknown, underlining the importance of future research. Yet, because competition is expected when and where shared resources are limited (61), for example floral resources in impoverished agricultural landscapes (13), it is remarkable that the addition of up to 35 honeybee hives had measurable effects on bumblebees in Irish heathlands. This highlights the need to consider competition, not only when beekeeping occurs in intensively farmed, flower-impoverished, landscapes, but also when it occurs in flower-rich environments, even within the native range of honeybees. Our results thereby have implications for the practice of heather honey production in heathlands both in Ireland and abroad.

Protecting wild bumblebee populations while also maintaining productive managed honeybee populations will require land managers and conservationists to work closely with beekeepers to evaluate appropriate stocking densities of honeybee hives in heathlands, especially near protected areas. To guide beekeepers and policy makers, it is important to understand how the carrying capacity of these heathlands is altered by the addition of honeybee hives, which will require further research.

## Author contribution statement

Conceptualization; KB, DS. Experimental design: KB, DS LH. Data collection: KB. Data analysis LH. Writing: KB DS LH.

## Data availability

All data and code is deposited on Figshare (10.6084/m9.figshare.28553870).

## Acknowledgements

We thank the beekeepers who shared the locations of their apiaries with us, the National Parks and Wildlife Service staff who helped us identify suitable sampling sites, and the landowners and Coillte staff who allowed us to survey on their land. We also thank Beau Williams for his field assistance, as well as Arrian Karbassioon and Linzi Thompson. Funding for this project was provided through the Eva Crane Trust (ECTA_20170910_Burns), an Irish Research Council Postgraduate Scholarship, in partnership with the Environmental Protection Agency (Project ID: GOIPG/2018/3411), and the Swedish Research Council Formas (Grant Number: 2018-01466).

## References

1. Potts SG, Biesmeijer JC, Kremen C, Neumann P, Schweiger O, Kunin WE. Global pollinator declines: trends, impacts and drivers. Trends in Ecology & Evolution. 2010;25(6):345–53.

2. Vanbergen AJ, Initiative tIP. Threats to an ecosystem service: pressures on pollinators. Frontiers in Ecology and the Environment. 2013;11(5):251–9.

3. IPBES. The assessment report of the Intergovernmental Science-Policy Platform on Biodiversity and Ecosystem Services on pollinators, pollination and food production. Bonn, Germany; 2016.

4. Mallinger RE, Gaines-Day HR, Gratton C. Do managed bees have negative effects on wild bees?: A systematic review of the literature. PLOS ONE. 2017;12(12):e0189268.

5. Paini DR. Impact of the introduced honey bee (Apis mellifera) (Hymenoptera: Apidae) on native bees: A review. Austral Ecology. 2004;29(4):399–407.

6. Goulson D. Effects of introduced bees on native ecosystems. Annual Review of Ecology, Evolution, and Systematics. 2003;34(Volume 34, 2003):1–26.

7. Furst MA, McMahon DP, Osborne JL, Paxton RJ, Brown MJF. Disease associations between honeybees and bumblebees as a threat to wild pollinators. Nature. 2014;506(7488):364–6.

8. Manley R, Boots M, Wilfert L. Emerging viral disease risk to pollinating insects: ecological, evolutionary and anthropogenic factors. Journal of Applied Ecology. 2015;52(2):331–40.

9. FAO. Crops and Livestock products. Live animals - bees. 2024;https://www.fao.org/faostat/(Accessed on 23 January 2025):Licence: CC-BY-4.0.

10. Torne-Noguera A, Rodrigo A, Osorio S, Bosch J. Collateral effects of beekeeping: Impacts on pollen-nectar resources and wild bee communities. Basic and Applied Ecology. 2016;17(3):199–209.

11. Page ML, Williams NM. Honey bee introductions displace native bees and decrease pollination of a native wildflower. Ecology. 2023;104(2):e3939.

12. Iwasaki JM, Hogendoorn K. Mounting evidence that managed and introduced bees have negative impacts on wild bees: an updated review. Current Research in Insect Science. 2022;2:100043.

13. Herbertsson L, Lindström SAM, Rundlöf M, Bommarco R, Smith HG. Competition between managed honeybees and wild bumblebees depends on landscape context. Basic and Applied Ecology. 2016;17(7).

14. Walther-Hellwig K, Fokul G, Frankl R, Büchler R, Ekschmitt K, Wolters V. Increased density of honeybee colonies affects foraging bumblebees. Apidologie. 2006;37(5):517–32.

15. Thomson D. Competitive interactions between the invasive European Honey Bee and native bumble bees. Ecology. 2004;85(2):458–70.

16. Goulson D, Sparrow K. Evidence for competition between honeybees and bumblebees; effects on bumblebee worker size. Journal of Insect Conservation. 2009;13(2):177–81.

17. Sutcliffe GH, Plowright RC. The effects of food supply on adult size in the bumble bee Bombus terricola Kirby (Hymenoptera: Apidae). The Canadian Entomologist. 1988;120(12):1051–8.

18. Ribeiro MF. Growth in bumble bee larvae: relation between development time, mass, and amount of pollen ingested. Canadian Journal of Zoology. 1994;72(11):1978–85.

19. Goulson D. Bumblebees: Behaviour and Ecology: Oxford University Press; 2003.

20. Greenleaf SS, Williams NM, Winfree R, Kremen C. Bee foraging ranges and their relationship to body size. Oecologia. 2007;153(3):589–96.

21. Lindström S, Smith HG. Konkurrens mellan honungsbin och vilda bin – evidens, kunskapsluckor och möjliga åtgärder. CEC Rapport Nr 6. Centrum för miljö-och klimatvetenskap, Lunds universitet. (English: Competition between honeybees and wild bees - evidence, knowledge gaps and potential mitigation strategies. CEC Report No. 6. Center for Environmental and Climate Science, Lund University). 2022.

22. Ruttner F. Biogeography and taxonomy of honeybees: Springer-Verlag Berlin Heidelberg; 1988.

23. Kryger P, Dupont YL. Konkurrence Mellem Honningbier og Vilde Bier. DCA – Nationalt Center for Fødevarer og Jordbrug. (English: Competition between honeybees and wild bees. DCA - National Center for Food and Agriculture). 2018.

24. Steffan-Dewenter I, Tscharntke T. Resource overlap and possible competition between honey bees and wild bees in central Europe. Oecologia. 2000;122(2):288–96.

25. Hudewenz A, Klein A-M. Competition between honey bees and wild bees and the role of nesting resources in a nature reserve. Journal of Insect Conservation. 2013;17(6):1275–83.

26. Franklin E, Carroll T, Blake D, Rickard K, Diaz A. Bumble bee forager abundance on lowland heaths is predicated by specific floral availability rather than the presence of honey bee foragers: evidence for forage resource partitioning. 2018. 2018;24.

27. Henry M, Rodet G. Controlling the impact of the managed honeybee on wild bees in protected areas. Scientific Reports. 2018;8(1):9308.

28. Magrach A, González-Varo JP, Boiffier M, Vilà M, Bartomeus I. Honeybee spillover reshuffles pollinator diets and affects plant reproductive success. Nature Ecology & Evolution. 2017;1(9):1299–307.

29. Lindström SAM, Herbertsson L, Rundlöf M, Smith HG, Bommarco R. Large-scale pollination experiment demonstrates the importance of insect pollination in winter oilseed rape. Oecologia. 2016;180(3):759–69.

30. Moquet L, Vanderplanck M, Moerman R, Quinet M, Roger N, Michez D, et al. Bumblebees depend on ericaceous species to survive in temperate heathlands. Insect Conservation and Diversity. 2017;10(1):78–93.

31. Descamps C, Moquet L, Migon M, Jacquemart AL. Diversity of the insect visitors on *Calluna vulgaris* (Ericaceae) in southern France heathlands. Journal of insect science (Online). 2015;15(1):130.

32. Koch H, Woodward J, Langat MK, Brown MJF, Stevenson PC. Flagellum removal by a nectar metabolite inhibits infectivity of a bumblebee parasite. Current Biology. 2019;29(20):3494–500.e5.

33. Kavanagh S, Gunnoo J, Marques Passos T, Stout JC, White B. Physicochemical properties and phenolic content of honey from different floral origins and from rural versus urban landscapes. Food Chemistry. 2019;272:66–75.

34. Forup ML, Memmott J. The relationship between the abundances of bumblebees and honeybees in a native habitat. Ecological Entomology. 2005;30(1):47–57.

35. Roubik DW, Buchmann SL. Nectar selection by Melipona and Apis mellifera (Hymenoptera: Apidae) and the ecology of nectar intake by bee colonies in a tropical forest. Oecologia. 1984;61(1):1–10.

36. Thomson JD. Pollen transport and deposition by Bumble Bees in Erythronium: Influences of floral nectar and bee grooming. Journal of Ecology. 1986;74(2):329–41.

37. Prado SG, Collazo JA, Marand MH, Irwin RE. The influence of floral resources and microclimate on pollinator visitation in an agro-ecosystem. Agriculture, Ecosystems & Environment. 2021;307:107196.

38. Department of Arts Heritage and the Gaeltacht. Site Synopsis - Wicklow Mountains SAC. National Parks and Wildlife Service 2017.

39. National Parks and Wildlife Service. Wicklow Mountains SAC (Site Code 002122) Conservation Objectives Supporting Document - Blanket Bogs and Associated Habitats. 2017.

40. EPA. Corine Land Cover 2018 Ireland. Environmental Protection Agency & European Environment Agency; 2018.

41. QGIS.org. QGIS Geographic Information System. Open Source Geospatial Foundation Project. http://qgis.org. 2020.

42. R Core Team. R: A language and environment for statistical computing.: R Foundation for Statistical Computing, Vienna, Austria. URL http://www.R-project.org; 2024.

43. Brooks ME, Kristensen K, van Benthem KJ, Magnusson A, Berg CW, Nielsen A, et al. glmmTMB balances speed and flexibility among packages for zero-inflated generalized linear mixed modeling. R Journal. 2017;9:378–400.

44. Hartig F. Residual diagnostics for hierarchical (Multi-Level / Mixed) regression models. Comprehensive R Archive Network. 2022.

45. Lenth R. emmeans: Estimated marginal means, aka least-squares means. R package version 1.5.2-12020.

46. Taillte Eireann. National Landcover Map for the Republic of Ireland. 2018.

47. Webb NR, Vermaat AH. Changes in vegetational diversity on remnant heathland fragments. Biological Conservation. 1990;53(4):253–64.

48. Willmer P. Pollination and floral ecology: Princeton University Press; 2011.

49. Spaethe J, Weidenmüller A. Size variation and foraging rate in bumblebees (*Bombus terrestris*). Insectes Sociaux. 2002;49(2):142–6.

50. Cresswell JE, Osborne JL, Goulson D. An economic model of the limits to foraging range in central place foragers with numerical solutions for bumblebees. Ecological Entomology. 2000;25(3):249–55.

51. Williams NM, Regetz J, Kremen C. Landscape-scale resources promote colony growth but not reproductive performance of bumble bees. Ecology. 2012;93(5):1049–58.

52. Goulson D, Peat J, Stout JC, Tucker J, Darvill B, Derwent LC, et al. Can alloethism in workers of the bumblebee, Bombus terrestris, be explained in terms of foraging efficiency? Animal Behaviour. 2002;64(1):123–30.

53. Beekman M, van Stratum P, Lingeman R. Diapause survival and post-diapause performance in bumblebee queens (Bombus terrestris). Entomologia Experimentalis et Applicata. 1998;89(3):207–14.

54. Owen RE. Body size variation and optimal body size of bumble bee queens (Hymenoptera: Apidae). The Canadian Entomologist. 1988;120(1):19–27.

55. Bommarco R, Lindström SAM, Raderschall CA, Gagic V, Lundin O. Flower strips enhance abundance of bumble bee queens and males in landscapes with few honey bee hives. Biological Conservation. 2021;263:109363.

56. Knee WJ, Medler JT. The seasonal size Increase of bumblebee workers (Hymenoptera: Bombus). The Canadian Entomologist. 1965;97(11):1149–55.

57. Plowright RC, Jay SC. Caste differentiation in bumblebees (Bombus Latr.: Hym.) I. — The determination of female size. Insectes Sociaux. 1968;15(2):171–92.

58. Shpigler H, Tamarkin M, Gruber Y, Poleg M, Siegel AJ, Bloch G. Social influences on body size and developmental time in the bumblebee Bombus terrestris. Behavioral Ecology and Sociobiology. 2013;67(10):1601–12.

59. Beekman M, Lew JB. Foraging in honeybees—when does it pay to dance? Behavioral Ecology. 2007;19(2):255–61.

60. Beekman M, Ratnieks FLW. Long-range foraging by the honey-bee, *Apis mellifera* L. Functional Ecology. 2000;14(4):490–6.

61. Birch LC. The meanings of competition. The American Naturalist. 1957;91(856):5–18.

