## Supplementary material for "Honeybee hive density has consequences for foraging bumblebees in Irish heathlands"

### SUPPLEMENTARY INFORMATION

Katherine L.W. Burns[1,2], Lina Herbertsson[3*], Dara A. Stanley[1,2*]

[1] School of Agriculture and Food Science, University College Dublin, Belfield, Dublin 2, Ireland

[2] Earth Institute, University College Dublin, Belfield, Dublin 2, Ireland

[3] Department of Biology, Lund University, SE-223 61 Lund, Sweden

*these authors contributed equally to the manuscript

### Figure S1

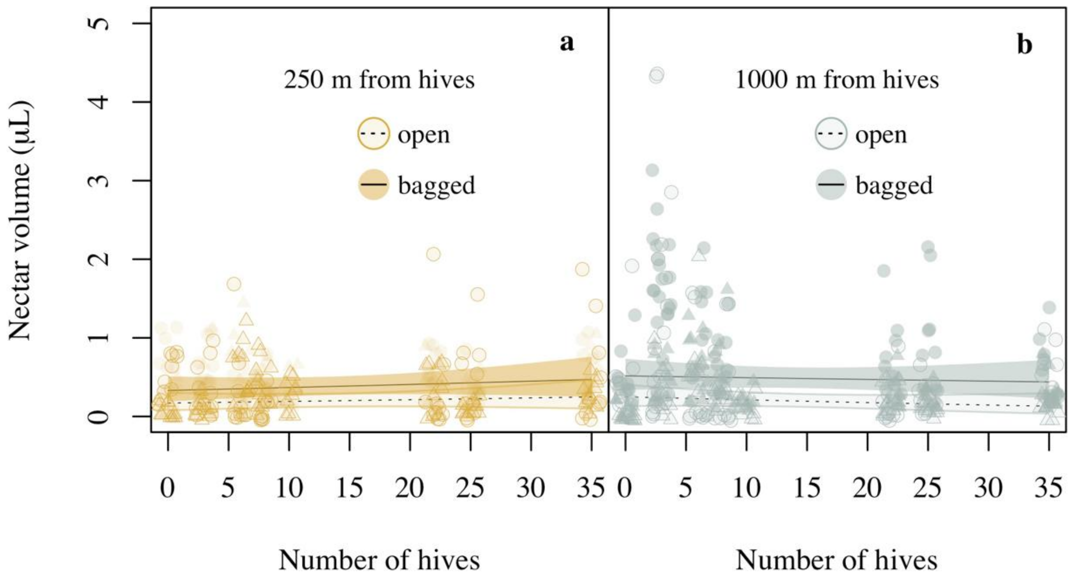


**Figure S1** The figure shows sampled nectar volumes from open (semi-open triangles/circles, dashed line) and bagged (filled circles/triangles, solid line) ling heather (*C. vulgaris*) **a)** 250 m from the hives and **b)** 1000 m from the hives. The interaction between treatment (bagged or open), hive density and distance was not significant. The figure shows model predictions with 95% confidence intervals. Raw data are displayed as open circles (first survey round) and triangles (second survey round). Raw data points have been jittered to reduce overlap and increase their visibility.

**Text S1 Influence of heathland**

The proportion of heathland had no influence on honeybee abundance (n = 32, heathland × hive density: $\chi^2_1$ = 0.11, p = 0.74, heathland: $\chi^2_1$ = 0.87, p = 0.35), bumblebee worker activity (n = 131, heathland × hive density: $\chi^2_1$ = 0.60, p = 0.44, heathland: $\chi^2_1$ = 0.06, p = 0.80) or proportional flowering heather (n = 160, heathland × hive density: $\chi^2_1$ = 0.72, p = 0.40, heathland: $\chi^2_1$ = 0.08, p = 0.78). Bumblebee abundance decreased with increasing proportion of heathland (n = 32, $\chi^2_1$ = 6.45, p = 0.01), and this was unrelated to hive density (heathland × hive density, n = 32, $\chi^2_1$ = 0.32, p = 0.57). The number of flowers visited by an individual of *B. lucorum* was unrelated to the proportion of heathland (n = 131, heathland × hive density: $\chi^2_1$ = 0.60, p = 0.44, heathland: $\chi^2_1$ = 0.06, p = 0.80). Thorax width was related to an interaction between hive density and proportion of heathland (n = 306, $\chi^2_1$ = 6.54, p = 0.01, Figure 7); a generally positive influence of heathland was counteracted by increasing hive density. When removing data points from the missing sheet, the full model did not converge and we removed species as a factor. To reduce unexplained variation, we then removed 92 measurements for *B. jonellus*. In this model, neither the interaction nor the main term effect of heathland were significant (n = 183, heathland × hive density: $\chi^2_1$ = 1.09, p = 0.30, heathland: $\chi^2_1$ = 2.19, p = 0.14), but width declined with hive density (n = 183, $\chi^2_1$ = 4.30, p = 0.04) and was smaller during the second round (n = 183, $\chi^2_1$ = 37.95, p < 0.0001). Nectar volumes were related to a significant interaction between treatment (open or bagged), hive density and proportion of heathland (n = 620, $\chi^2_1$ = 3.85, p = 0.05, Figure S2). This was driven by an interaction between hive density and proportion of heathland for bagged (contrast: $F_{1,607}$ = 4.18, p = 0.04), but not for open flowers ($F_{1,607}$ = 0.08, p = 0.78); nectar volumes of bagged flowers decreased with hive density when the proportion of heathland was low (Figure S2a) and increased when the proportion of heathland was high (Figure S2b). Nectar volumes of open flowers were lower and unrelated to heathland ($F_{1,607}$ = 2.08, p = 0.09) as well as hive density ($F_{1,607}$ = 0.38, p = 0.53).

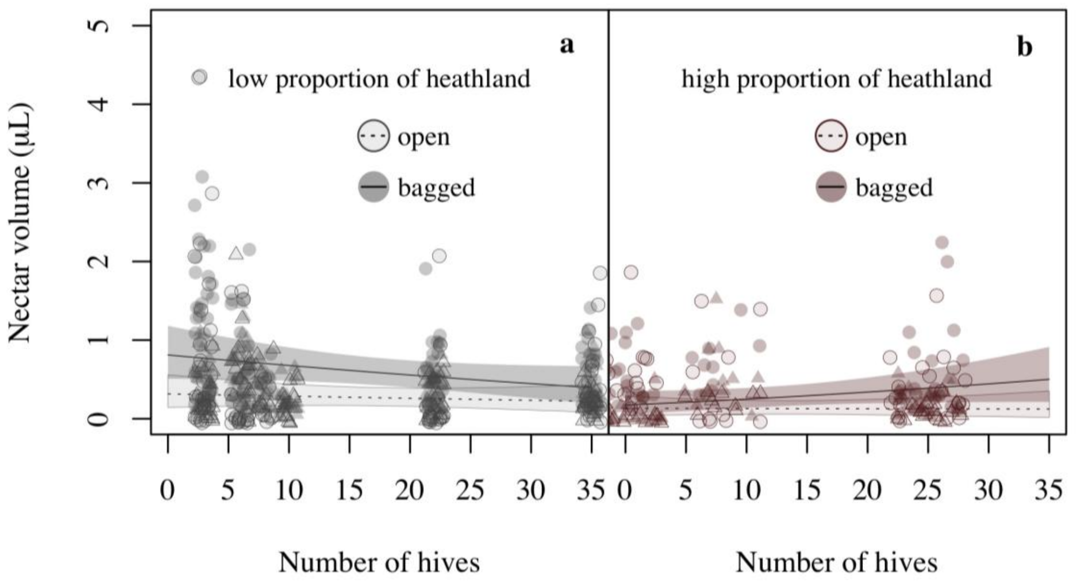


**Figure S2** The figure shows sampled nectar volumes from open (semi-open circles/triangles, dashed line) and bagged (filled circles/triangles, solid line) ling heather (*C. vulgaris*) **a)** where the proportion of heathland was lower than average (57%) and **b)** where it was higher than the average (57%). The figure shows model predictions with 95% confidence intervals for a) the lowest and b) highest proportion of heathland. Raw data are displayed as open circles (first survey round) and triangles (second survey round). Raw data points have been jittered to reduce overlap and increase their visibility.

**Table S1.** Total species and abundances of bees (*A. mellifera, B. jonellus, B. lucorum agg.*, *B. monticola, B. pascuorum,* and *B. pratorum*) summed across all 250 m sites (250 m from nearest apiary) and 1000 m sites (1000 m from nearest apiary).

| **Distance** | **Bee species** | **Species abundance** |
|---|---|---|
| 250 m | *A. mellifera* | 65 |
| | *B. jonellus* | 7 |
| | *B. lucorum agg.* | 52 |
| | *B. pascuorum* | 4 |
| | **Total** | 128 |
| 1000 m | *A. mellifera* | 117 |
| | *B. jonellus* | 18 |
| | *B. lucorum agg.* | 58 |
| | *B. monticola* | 2 |
| | *B. pascuorum* | 2 |
| | *B. pratorum* | 1 |
| | **Total** | 199 |

**Table S2.** Total abundance of each species (*A. mellifera*, *B. jonellus*, *B. lucorum agg.*, *B. monticola*, *B. pascuorum*, and *B. pratorum*) observed foraging on available floral resources (*C. vulgaris*, *Erica cinerea*, *Erica tetralix*, and *Ulex europaeus*) during transect observations.

| | **Forage species** | | | | |
|---|---|---|---|---|---|
| **Bee species** | ***C. vulgaris*** | ***E. cinerea*** | ***E. tetralix*** | ***U. europaeus*** | **Total** |
| *A. mellifera* | 174 | 5 | 0 | 3 | 182 |
| *B. jonellus* | 23 | 2 | 0 | 0 | 25 |
| *B. lucorum agg.* | 109 | 1 | 1 | 0 | 111 |
| *B. monticola* | 2 | 0 | 0 | 0 | 2 |
| *B. pascuorum* | 2 | 1 | 0 | 0 | 6 |
| *B. pratorum* | 1 | 0 | 0 | 0 | 1 |
| **Total** | 311 | 12 | 1 | 3 | 327 |
